# QTL mapping of natural variation reveals that the developmental regulator *bruno* reduces tolerance to *P*-element transposition in the *Drosophila* female germline

**DOI:** 10.1101/282541

**Authors:** Erin S. Kelleher, Jaweria Jaweria, Uchechukwu Akoma, Lily Ortega, Wenpei Tang

## Abstract

Transposable elements (TEs) are obligate genetic parasites that propagate in host genomes by replicating in germline nuclei, thereby ensuring transmission to offspring. This selfish replication not only produces deleterious mutations---in extreme cases, TE mobilization induces genotoxic stress that prohibits the production of viable gametes. Host genomes could reduce these fitness effects in two ways: resistance and tolerance. Resistance to TE propagation is enacted by germline specific small-RNA-mediated silencing pathways, such as the piRNA pathway, and is studied extensively. However, it remains entirely unknown whether host genomes may also evolve tolerance, by desensitizing gametogenesis to the harmful effects of TEs. In part, the absence of research on tolerance reflects a lack of opportunity, as small-RNA-mediated silencing evolves rapidly after a new TE invades, thereby masking existing variation in tolerance.

We have exploited the recent the historical invasion of the *Drosophila melanogaster* genome by *P*-element DNA transposons in order to study tolerance of TE activity. In the absence of piRNA-mediated silencing, the genotoxic stress imposed by *P*-elements disrupts oogenesis, and in extreme cases leads to atrophied ovaries that completely lack germline cells. By performing QTL-mapping on a panel of recombinant inbred lines (RILs) that lack piRNA-mediated silencing of *P*-elements, we uncovered multiple QTL that are associated with differences in tolerance of oogenesis to *P*-element transposition. We localized the most significant QTL to a small 230 Kb euchromatic region, with the LOD peak occurring in the *bruno* locus, which codes for a critical and well-studied developmental regulator of oogenesis. We further demonstrate tolerant alleles are associated with reduced *bruno* expression, and that multiple *bruno* loss-of-function alleles are strong dominant suppressors of ovarian atrophy, allowing for the development of mature egg-chambers in the face of *P*-element activity. Genetic and cytological analyses suggest that *bruno* tolerance is explained by enhanced retention of germline stem cells in dysgenic ovaries, which are typically lost due to DNA damage. Our observations reveal segregating variation in TE tolerance for the first time, and implicate gametogenic regulators as a source of tolerant variants in natural populations.

## INTRODUCTION

Transposable elements (TEs) are omnipresent and abundant constituents of eukaryotic genomes, comprising up to 80% of genomic DNA in some lineages (reviewed in [1]). The evolutionary footprint of TEs is extensive, including dramatic genome size expansions [2,3], acquisition of new regulatory networks [4,5], structural mutations [6], novel genes [7–9], and adaptive insertions [10–12]. However, the charismatic and occasionally beneficial impact of TEs over evolutionary time masks their fundamental identity as intragenomic parasites and mutagens. In addition to causing deleterious mutations [13,14], TEs can exert lethal, genotoxic effects on host cells by producing abundant double-stranded breaks (DSBs) during insertion and excision [15,16]. TEs are therefore intragenomic parasites.

Host developmental and evolutionary responses to parasites, pathogens, and herbivores are broadly delineated into two categories: resistance and tolerance (reviewed in [17,18]). Mechanisms of resistance prevent–or limit the spread of–infection or herbivory. By contrast, mechanisms of tolerance do not affect propagation, but rather limit the fitness costs to the host. With respect to TEs, resistance by eukaryotic genomes is enacted by small RNA mediated silencing pathways [19], and <u>K</u>ruppel-<u>a</u>ssociated <u>b</u>ox <u>z</u>inc-<u>f</u>inger proteins (KRAB-ZFPs)[20], which regulate the transcription and subsequent transposition of endogenous TEs. However, it remains unknown whether genomes can also evolve tolerance of TEs, by altering how host cells are affected by TE activity. Tolerance therefore represents a wholly unexplored arena of the evolutionary dynamics between TEs and their hosts. Germline tolerance of TEs is predicted to be of particular importance, due to the significance of this cell lineage in ensuring the vertical transmission of the parasite, and the reproductive fitness of the host.

The absence of research on tolerance is at least partially due the primacy of resistance: endogenous TEs are overwhelmingly repressed by host factors in both germline and somatic tissues [19,21,22]. However, the invasion of the host genome by a novel TE family, which happens recurrently over evolutionary time scales (reviewed in [23]), provides a window of opportunity through which tolerance could be viewed, both empirically and by natural selection. The absence of evolved resistance in the host against a new invader could reveal differential responses of germline cells to unrestricted transposition. A classic example of genome invasion by a novel TE is provided by *P*-elements, DNA transposons that have recently colonized two *Drosophila* species. *P*-elements first appeared in genomes of *D. melanogaster* around 1950 (reviewed in [24]), and later colonized its sister species *D. simulans* around 2006 [25,26]. Particularly for *D. melanogaster*, a large number of naïve strains collected prior to *P*-element invasion are preserved in stock centers and laboratories, providing a potential record of ancestral genetic variation in tolerance [27,28]. Furthermore, 15 of these naïve strains were recently used to develop the *Drosophila* Synthetic Population Resource (DSPR), a panel of highly Recombinant Inbred Lines (RILs) that serve as a powerful toolkit for discovering the natural genetic variants influencing quantitative traits [29–31].

Here, we harness the mapping power of the DSPR to screen for phenotypic and genetic variation in the tolerance of the *D. melanogaster* female germline to unrestricted *P*-element activity. We developed a novel screen for phenotypic variation in host tolerance by taking advantage of the classic genetic phenomenon of hybrid dysgenesis, in which TE families that are inherited exclusively paternally can induce a sterility syndrome in offspring, due to an absence of complementary maternally-transmitted regulatory small RNAs (piRNAs, reviewed in [24,32,33]). The dysgenesis syndrome induced by *P*-elements in the female germline is particularly severe, and can be associated with a complete loss of germline cells [15,34,35]. *P*-element hybrid dysgenesis is directly related to genotoxic stress, as apoptosis is observed in early oogenesis in dysgenic females [15], and the DNA damage response factors *checkpoint kinase 2* and *p53* act as genetic modifiers of germline loss [35]. Variation in the sensitivity of the DNA damage response to double-stranded breaks therefore represents one potential cellular mechanism for tolerance.

By phenotyping the germline development of >32,000 dysgenic female offspring of RIL mothers, we uncovered substantial heritable variation in female germline tolerance of *P*-element activity. We furthermore mapped this variation to a small 230 Kb quantitative trait locus (QTL) on the second chromosome, and associated it with the differential expression of *bruno*, a well-studied developmental regulator of oogenesis with no known function in TE repression or DNA damage response [36–39]. We further demonstrate that *bruno* loss-of function alleles act as dominant suppressors of germline loss, and relate these effects to the retention of germline stem cells (GSCs) in dysgenic females. Our findings represent the first demonstration of natural variation in TE tolerance in any organism. They further implicate regulators of gametogenesis such as *bruno*, as a source of tolerant variants that could be beneficial when new TEs invade the host.

## RESULTS

### Maternal genotype, but not zygotic age, is a strong predictor of dysgenic F1 atrophy

We first sought to uncover phenotypic variation in germline tolerance of *P*-element activity among a panel of highly recombinant inbred lines (RILs) derived from eight founder genomes [29]. To quantify tolerance, we performed dysgenic crosses between RIL females and males from the *P*-element containing strain Harwich (Fig 1A). Harwich is a strong paternal inducer of hybrid dysgenesis, producing F1 females with 100% atrophied ovaries in crosses with naïve females at the restrictive temperature of 29°C [28]. We therefore performed our crosses at 25°C, a partially permissive temperature at which intermediate levels of ovarian atrophy are observed [40,41]. Because *P*-element dysgenic females may recover their fertility as they age, through zygotic production of *P*-element derived piRNAs [42], we assayed both 3-day old and 21-day old F1 female offspring from each RIL. In total, we documented the incidence of atrophied ovaries among 17,150 3-day-old and 15,039 21-day-old F1 female offspring, and estimated the proportion of F1 atrophy within broods of ≥ 20 3-day old and 21-day old offspring from 592 and 492 RILs, respectively (Fig 1B, C). Notably, because we phenotyped F1 offspring, our phenotypic variation will be determined only by variants in which one of the RIL alleles is at least partially dominant to the Harwich allele, as well as any maternal effects determined solely by the RIL genotype.

**Fig 1.**
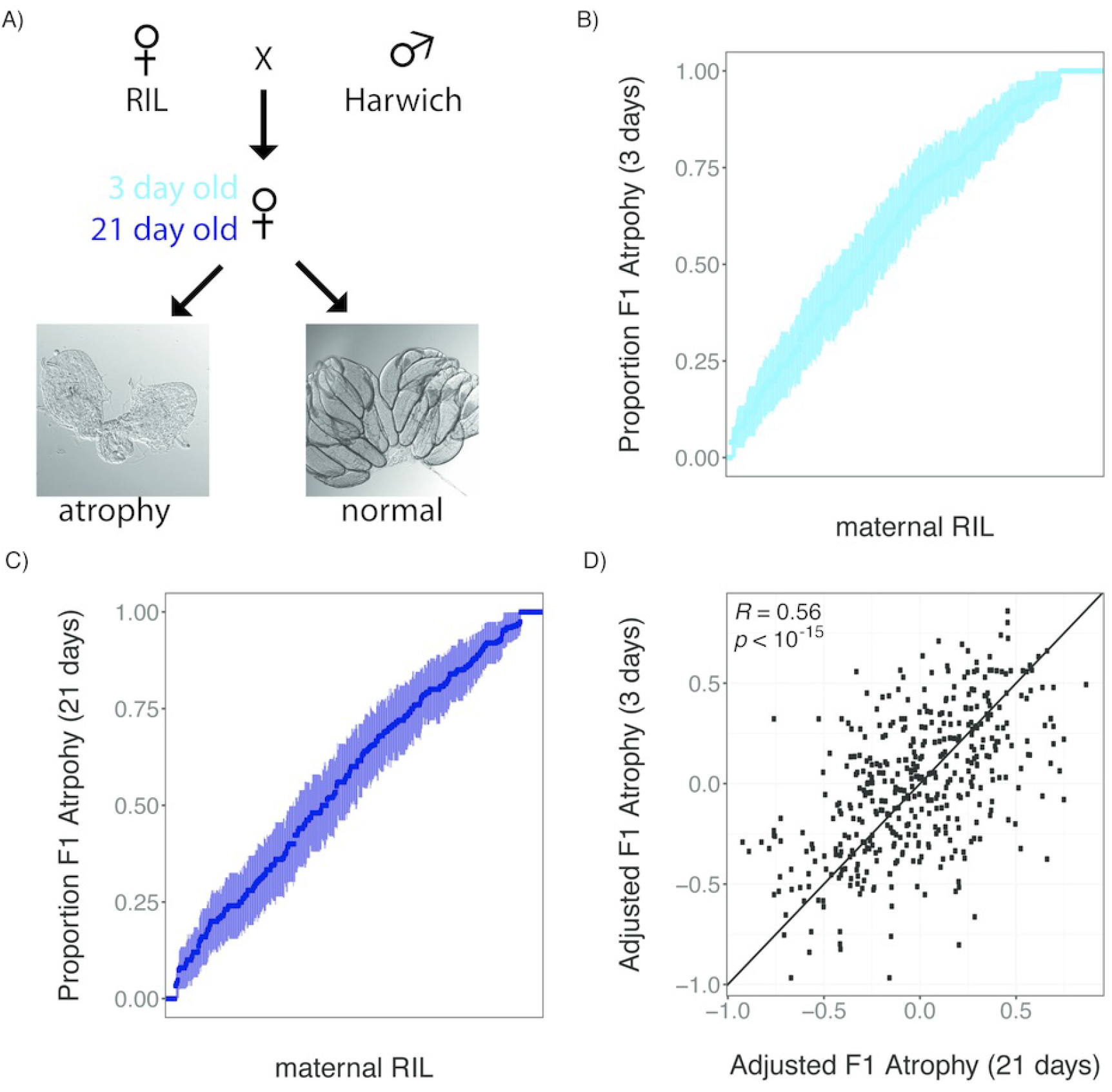
Heritable phenotypic variation in tolerance of *P*-element activity in the female germline. A) Crossing scheme for documenting variation in tolerance of *P*-element activity among RIL offspring. Representative images of atrophied and normal ovaries were published previously [35]. B and C) phenotypic variation in the proportion of 3-day old (B) and 21-day old (C) F1 female offspring of different RILs. RILs are sorted according the proportion of F1 atrophy observed in their offspring and error bars indicate the standard error of the estimated proportion. D) Scatterplot of the arsine-transformed proportion of F1 atrophy observed among 3-day old and 21-day old offspring of the same line, after accounting for the effects of experimenter and experimental block. The individual numerical values for panels B, C, and D can be found in S1 Data, S2 Data and S3 Data, respectively.

We observed continuous variation in the proportion of F1 atrophy among both 3-day old and 21-day old offspring of different RIL genotypes (Fig 1B, 1C). After accounting for the effects of experimenter and experimental block, the incidence of ovarian atrophy is strongly correlated among 427 RILs for which we sampled broods of both 3-day old and 21-day old F1 females (Pearson’s *R* = 0.56, *p >* 10^−15^, Fig 1D). Because broods of different age classes were sampled from separate crosses and experimental blocks, this correlation strongly implies that phenotypic differences are explained by the maternal genotype. Indeed, based on the F1 atrophy proportions measured in broods of different ages, we estimate that the broad-sense heritability of F1 ovarian atrophy among the RIL offspring was 40.35% (see Materials and Methods). Furthermore, we saw greater reproducibility across a small sample of 14 RILs for which we phenotyped two independent 21-day broods (Pearson’s *R =* 0.97, *p >* 10^−8^), suggesting even higher heritability among offspring of the same age class.

Despite the previous observation of developmental recovery from hybrid dysgenesis [42], the relationship between age and the proportion of F1 atrophy is only marginally significant in our data (*F_1,1037_*= 3.57, *p =* 0.058). Furthermore, 21-day old dysgenic females exhibited only a 0.63% decrease in the proportion of F1 atrophy when compared to 3-day old females, indicating that the overall effect of age in our crosses was very modest. Finally, we did not observe a group of RILs in which ovarian atrophy is much more common among 3-day-old as compared to 21-day-old F1 females (Fig 1D), as would be predicted if there were genetic variation for developmental recovery across the RIL panel. The absence of developmental recovery of in our experiments could reflect differences in developmental temperature between our two studies (22°C in [42] and 25°C here). Alternatively, the causative variant that allows for developmental recovery could be absent from the founder RILs.

### QTL Mapping of Dysgenic F1 Atrophy

To identify the genomic regions that harbor causative genetic variation in tolerance, we performed a QTL analysis using the published RIL genotypes [29]. In these data, the founder allele (A1-A8) carried by each RIL is inferred probabilistically for 10 Kb windows along the euchromatic regions of the major autosomes 2 and 3, and the X-chromosome [29]. The fourth chromosome is ignored because the absence of recombination makes it uninformative for QTL mapping (reviewed in [43]).

Consistent with the strongly correlated phenotypes of 3-day-old and 21-day-old offspring (Fig 1D), we identified a single major effect QTL associated with phenotypic variation at both developmental time points (Fig 2A). Additionally, the Δ2-LOD drop confidence intervals (Δ2-LOD CI) of the LOD peak from each analysis are both narrow (<300 Kb) and highly overlapping (Table 1). The peak explains 14.2% and 14.8% of variation in ovarian atrophy among 3-day-old and 21-day-old F1 females, indicating it is a major determinant of heritable variation. Multiple minor peaks close to the centromere of 3L may represent another source of heritable variation in 3-day-old females.

**Fig 2.**
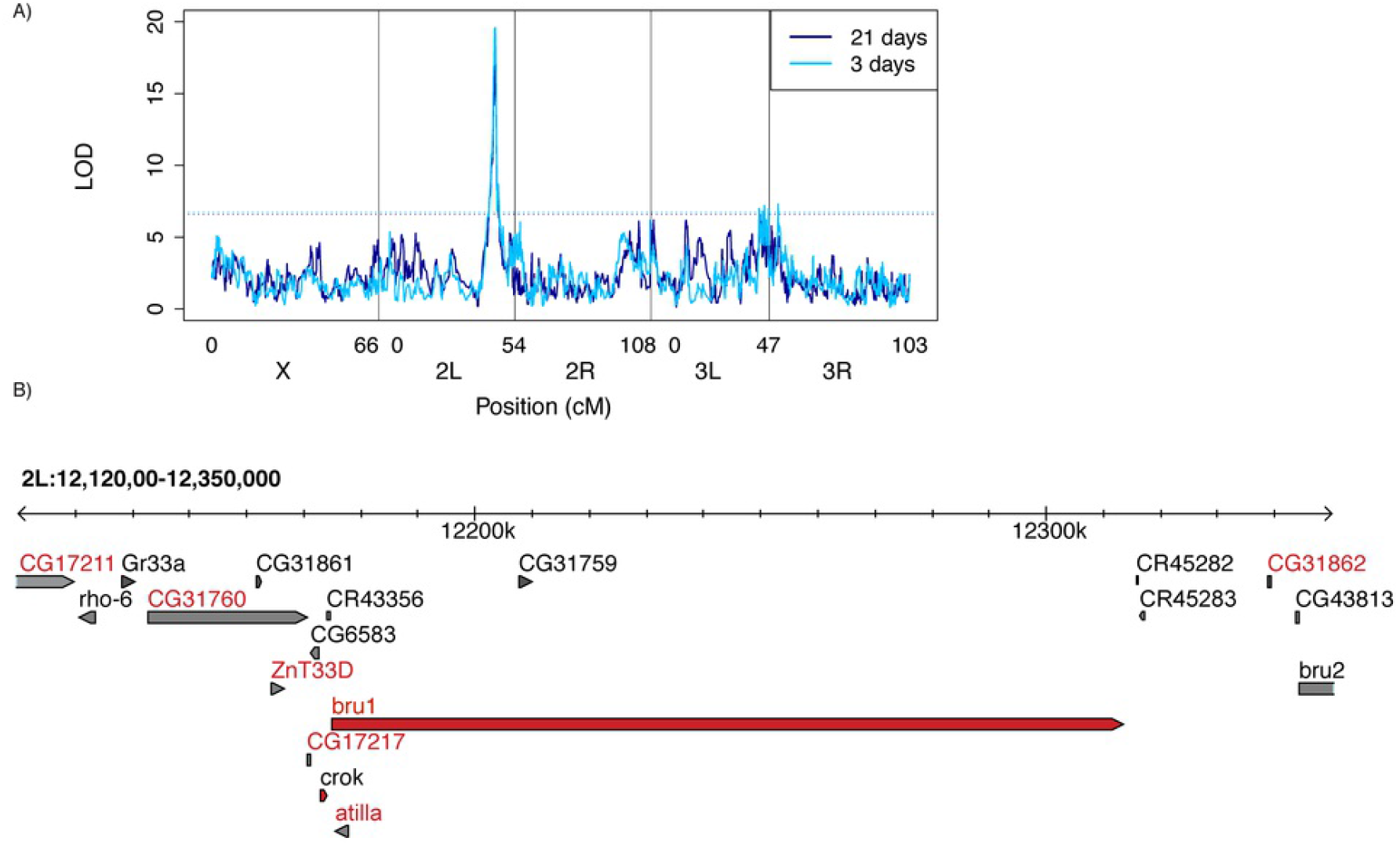
QTL mapping of variation in *P*-element tolerance. A) <u>L</u>ogarithm of the <u>o</u>dds <u>r</u>atio (LOD) of the observed association between maternal RIL genotype [29], and the adjusted proportion of F1 atrophy phenotype. Higher LOD scores correspond to stronger evidence of linkage, and significant LOD scores are above the threshold (dotted line), which was obtained from 1,000 permutations of the observed data. B) 2 LOD-drop confidence interval of the QTL peak based on a combined QTL analysis including both 3-day old and 21-day old F1 females. Genes indicated by red letters are potentially effected by polymorphisms in the RIL founders that are in phase the inferred allelic classes (Fig 3 [44]). Gene models indicated in red are highly expressed in the *D. melanogaster* ovary [45]. The individual numerical values required to generate LOD plots for 3-day old and 21-day old F1 females can be found in S4 Data and S5 Data, respectively.

**Table 1.** QTL peak positions. The position of the major QTL peak for 3-day old F1 females, 21-day old F1 females, and the combined data set are shown. For each analysis, the peak position, Δ2-LOD drop Confidence Interval, and Bayesian credible interval [46] in *Drosophila melanogaster* genome release 6 (dm6 [47]) are provided. The percent of phenotypic variation explained by the QTL peak is based on the genotype of each sampled RIL at the LOD peak position. The individual numerical values required to identify LOD peaks and intervals for 3-day, 21-day old, and both F1 females can be found in S4 Data, S5 Data, S3 Data respectively.

| Data Set | LOD Score | Peak Position | $\Delta 2$-LOD CI | Bayes CI | % variation |
| --- | --- | --- | --- | --- | --- |
| 3-days | 19.59 | 2L:12260000 | 2L:12080000-12380000 | 2L:12080000-12320000 | 14.2 |
| 21-days | 17.22 | 2L:12250000 | 2L:12110000-12390000 | 2L:12160000-12330000 | 14.8 |
| both | 34.90 | 2L:12260000 | 2L:12120000-12350000 | 2L:12180000-12310000 | 13.8 |

To further narrow the location of genetic variation in tolerance, we took advantage of the striking concordance in QTL mapping for the 3-day old and 21-day old datasets, and performed a combined analysis including all 660 RILs whose F1 offspring were sampled at either developmental time point. F1 female age was included as a covariate (see methods). From this analysis we obtained a final Δ2-LOD CI for the major QTL peak, which corresponds to a 230 Kb genomic region containing 18 transcribed genes, 15 protein-coding and 3 non-coding (Fig 2B). Simulation testing of the statistical properties of the DSPR indicate that a causative variant explaining 10% of phenotypic variation lies within the Δ2-LOD drop CI 96% of time for sample sizes of 600 RILs [30]. Furthermore, overestimation of variance explained by a QTL (*i.e*. the Beavis effect), is rare in DSPR studies sampling greater than 500 RILs, particularly for variants explaining ≥10% of variation [48]. Therefore, given our sample (660 RILs) and effect (~14%) sizes, the Δ2-LOD CI we infer for our major peak should be conservative. Indeed, Bayesian credible intervals, an alternate approach for identifying QTL windows [46], are even narrower than those estimated by Δ2-LOD CI (Table 1).

Of the 18 genes within the QTL peak, only *bruno* and *crooked* are highly expressed in the *D. melanogster* ovary [45]. While *bruno* is translational repressor whose major essential role is in oogenesis [37,49,50], *crooked* is a more broadly expressed component of septate junctions, which is essential for viability [51]. The LOD peak resides within the 138 Kb *bruno* locus which, in addition to its function, make it the strongest candidate for the source of causative variation.

### Two classes of tolerance alleles

We next sought to partition founder alleles at the QTL peak into phenotypic classes, in order to better understand the complexity of causative genetic variation. First we identified all sampled RILs whose genotype was probabilistically assigned (*p* > 0.95) to a single founder (A1-A8) at the LOD peak, and estimated the phenotypic effect associated with each founder allele (Fig 3A, 3B). We then used stepwise regression (see Materials and Methods) to identify the minimum number of allelic classes required to explain phenotypic variation associated with the founder alleles. For both age classes, we found strong evidence for 2-allelic classes, one sensitive and one tolerant, which were sufficient explain phenotypic variation in tolerance. This implies that the major QTL peak could correspond to a single segregating genetic variant. Furthermore, with the exception of founder A8, founder alleles were assigned to the same allelic class for both age cohorts, revealing allelic behavior is highly biologically reproducible.

**Fig 3.**
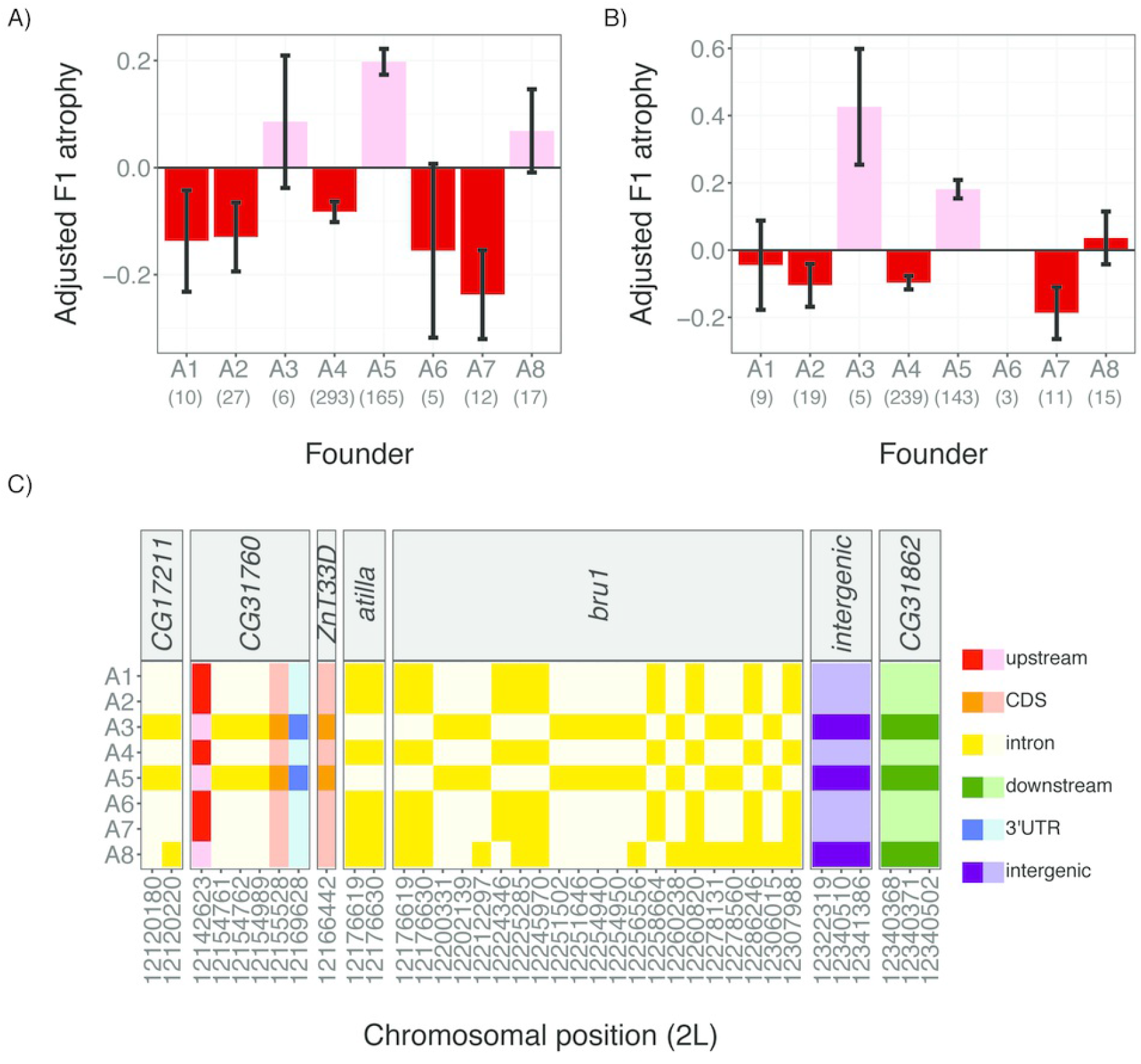
Phenotypic classes of founder alleles at the major QTL peak. The mean adjusted F1 atrophy among 3-day old females A) and 21-day old females B) is shown for RILs carrying each of the 8 founder alleles. Error bars denote standard error. QTL phasing (see methods) detects two allelic classes for both the 3 day-old and 21 day-old phenotypes: a sensitive allele that increases the odds F1 ovarian atrophy (pink) and a tolerant allele that decreases the odds of F1 ovarian atrophy 1 (red). The assignment of founder alleles to phenotypic classes across ages is consistent for all founders except A8. C) In-phase SNPs in population A RIL founder genomes, indicated by their position in the *Drosophila melanogaster* reference genome dm6 [47]. SNPs are colored according to the function of the affected sequence, and shaded according to whether the founder exhibits the reference (light) or alternate (dark) allele. The individual numerical values required to generate bar plots for panels A and B can be found in S4 and S5 Data, respectively. In-phase polymorphisms represented in panel C are provided in S1 Table.

To further study phenotypic differences between tolerant and sensitive alleles, we identified three pairs of background-matched RILs, which exhibited a tolerant (founder A4) or a sensitive (founder A5) haplotype across the QTL window, but otherwise shared a maximal number of alleles from the same founder across the remainder of the genome. Consistent with our QTL mapping, these RIL pairs differed dramatically in the incidence of ovarian atrophy they displayed in crosses with Harwich males (Fig 4A). While we did not detect a significant effect of genetic background (drop-in-deviance = 3.57, *df*= 2, *p*= 0.17), the QTL haplotype was strongly associated with the incidence of ovarian atrophy (drop-in-deviance = 52.01, *df*= 1, *p* = 5.36 × 10^−13^). RILs carrying the tolerant haplotype exhibited 39% less F1 ovarian atrophy than those carrying the sensitive haplotype.

**Fig 4.**
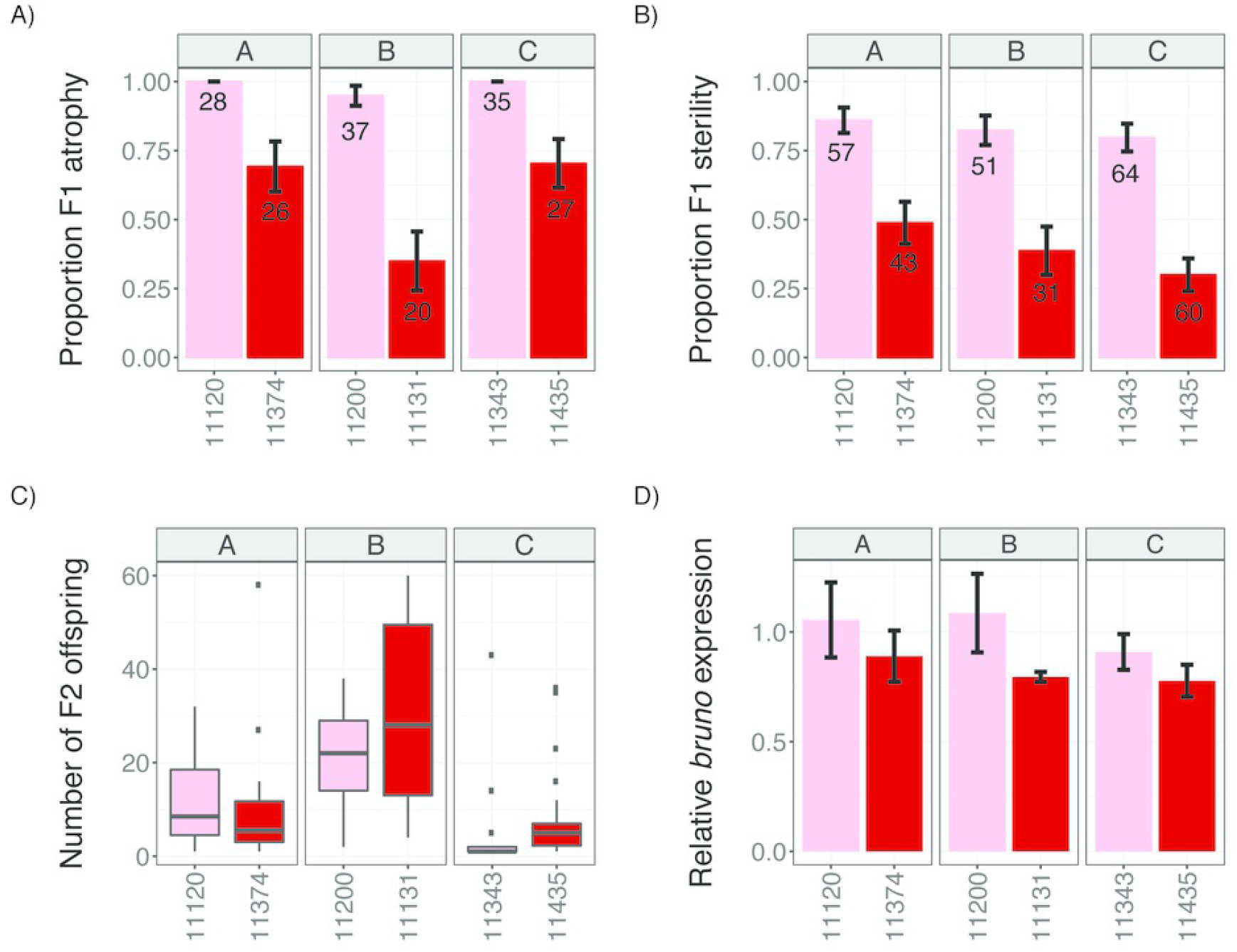
Tolerant alleles are associated with enhanced fertility and reduced *bruno* expression. Phenotypes of background-matched RILs (A, B and C) carrying sensitive (A5, pink) and tolerant (A4, red) haplotypes across the QTL peak are compared. A-B) Incidence of ovarian atrophy (A), and sterility (B) among dysgenic F1 female offspring of crosses between RIL females and Harwich males. Numbers in each bar indicate the sample size. C) Offspring production of fertile F1 females from B. D) Ovarian *bruno* expression relative to *rpl32* in each RIL. Error bars in A, B, and D indicate the standard error. The individual numerical values required to generate bar and box-plots for panels A, B/C, and D can be found in S6 Data, S7 Data, and S8 Data, respectively.

To determine whether reduced ovarian atrophy conferred by tolerant alleles increases female reproductive fitness, we examined the presence and number of F2 adults produced by young (0-5 day-old) F1 female offspring of tolerant and sensitive dysgenic crosses (Fig 4B). The proportion of F1 sterility was somewhat lower than the proportion of F1 atrophy for the same dysgenic cross (Fig 4A vs. 4B), consistent with loss of germline cells in early adult stages. Equivalent to ovarian atrophy, there was no significant effect of genetic background (drop-in-deviance = 5.26, *df*= 2, p= 0.07), but the QTL haplotype was strongly associated with F1 sterility (drop-in-deviance = 65.787, *df*= 1, *p* = 5.55 × 10^−16^). F1 females carrying the tolerant haplotype exhibited a 54% reduction in sterility as compared to those carrying the sensitive haplotype. Interestingly, when we examine the number of F2 offspring produced by fertile F1 females from resistant and tolerant crosses (Fig 4C), we detect dramatic effects of genetic background (*F_2,110_* = 29.05, *p* = 7.48 × 10^−11^), but no significant effect of the tolerant allele (*F_1,110_* = 1.01, *p* = 0.31). Therefore, while tolerant alleles enhance female reproductive fitness by increasing the odds of fertility, other genetic factors likely determine the number of offspring produced by those fertile females.

### Tolerant alleles are associated with reduced *bruno* expression

In light of the simple biallelic behavior of our phenotype, we sought to identify polymorphisms within the founder strains whose genotypic differences matched their phenotypic classifications (*i.e*. “in-phase” polymorphisms [29,31], Fig 3C). We excluded A8 from these analyses due to the ambiguity of its allelic class. In total we identified 36 in-phase single nucleotide polymorphisms (SNPs), which potentially affect the function of only seven transcribed genes in the QTL interval ([44] S1 Table, Fig 2B, 3C). We did not identify any in-phase, segregating TE insertions, although a recent reassembly of the founder A4 genome based on long single-molecule real-time sequencing reads suggests many TE insertions remain un-annotated [52]. Focusing on *bruno* and *crooked*, the two genes in the QTL window that are highly expressed in the ovary [45], 22 of the in-phase SNPs are within *bruno* introns, while none are found in the gene-body or upstream of *crooked*. Furthermore, none of the 36 in-phase SNPs are non-synonymous, implying a regulatory difference between the tolerant and sensitive alleles.

To determine if tolerance is associated *bruno* regulation, we compared ovarian expression of *bruno* in young (3 day-old) females from our background matched RIL pairs carrying a tolerant or sensitive haplotype across the QTL locus. While we observed only modest effects of genetic background on *bruno* expression (Likelihood Ratio Test = 6.57, *df*=2, *p* = 0.04), we observed dramatic effects of founder haplotype at the QTL window (Likelihood Ratio Test = 29.47, *df*=1, *p* = 5.67 × 10^−8^). Across genetic backgrounds, tolerant alleles were associated with a 20% reduction in *bruno* expression (95% CI: 14-26%), suggesting that *bruno* function reduces germline tolerance of *P*-element activity.

### *bruno* is a strong dominant suppressor of *P*-element induced ovarian atrophy

Given the differences in *bruno* expression between sensitive and tolerant alleles, we wondered whether *bruno* loss-of-function alleles affect the atrophy phenotype. For comparison, we also considered available alleles from three other ovary-expressed genes that are located within the Δ2-LOD CI of the 21-day old or 3-day old female analyses, but not in the combined analysis: *ced-12, Rab6* and *ThrRS*. We reasoned that since the causative variant is almost certainly not recessive, being found only in the maternal RIL genotype, mutant alleles might also exhibit non-recessive effects. We therefore used balanced heterozygous females as mothers in dysgenic crosses with Harwich males, and compared the incidence of F1 ovarian atrophy among their 3-7 day-old F1 females (*mutant/+ vs. balancer/+*). Strikingly, while *ced-12, Rab6* and *ThrRS* alleles had no effect on the incidence of ovarian atrophy (Fig 5A), two different *bruno* alleles (*bruno^RM^* and *bruno^QB^*) acted as dominant suppressors (Fig 5B). In contrast to their balancer control siblings, who exhibited 64-75% ovarian atrophy, *bruno^RM^/+* and *bruno^QB^/+* offspring exhibited only 13% and 20% atrophy, respectively, a dramatic reduction. We further observed that an independently-derived *bruno* deficiency [53,54] suppresses ovarian atrophy to a similar degree (Fig 5B), indicating these effects cannot be attributed to a shared linked variant on the *bruno^RM^* and *bruno^QB^* chromosomes [49].

**Fig 5.**
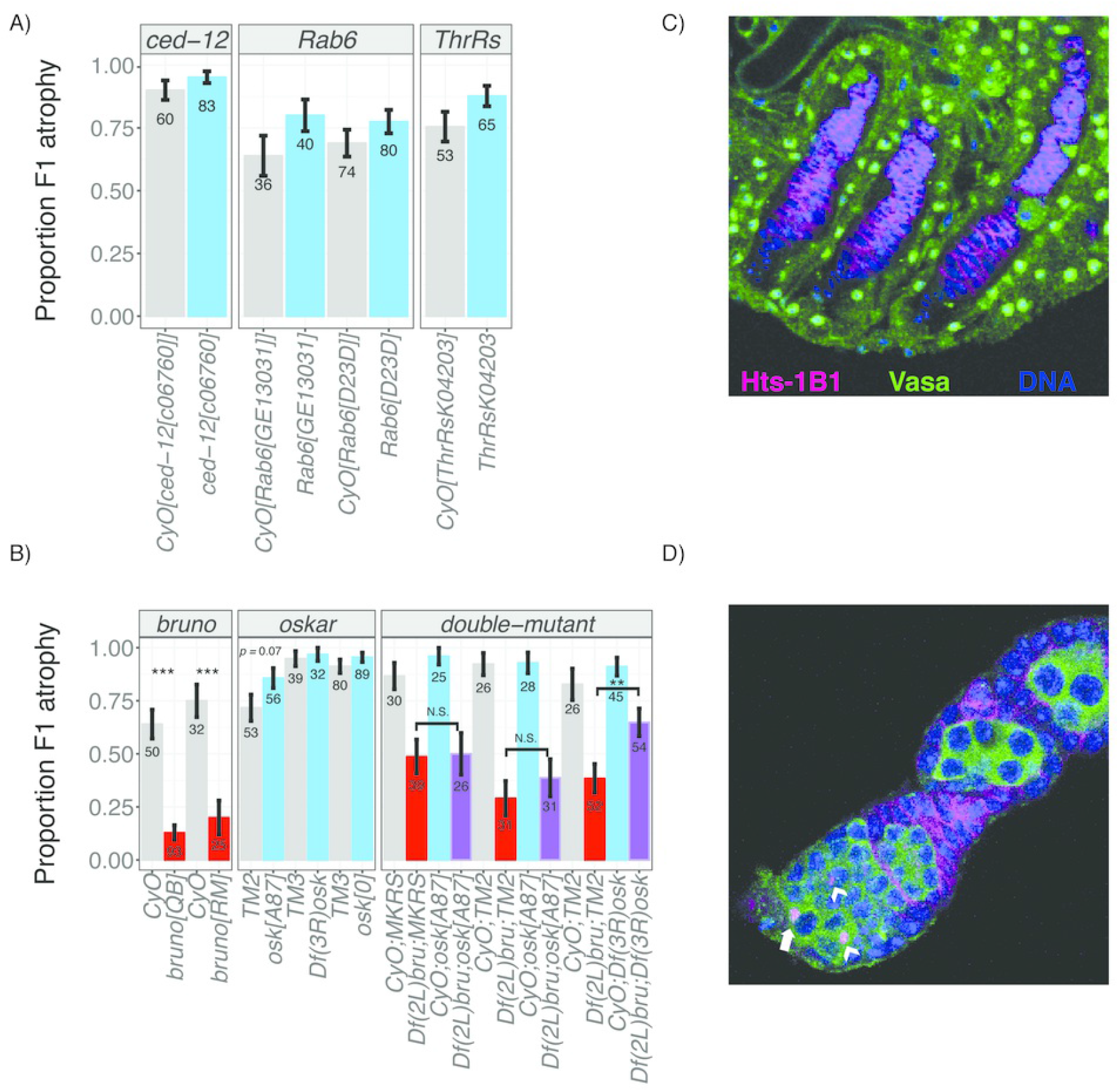
Mutational analysis of candidate genes. A) Loss-of-function heterozygotes for three candidate genes are compared to their siblings inheriting the balancer chromosome *CyO*, in order to detect zygotic effects on the incidence of ovarian atrophy among 3-7 day-old dysgenic F1 offspring. B) Single and double loss-of-function and deficiency heterozygotes for *bruno* and *oskar* are compared to their siblings inheriting the balancer chromosomes *CyO* (*bruno*) and *TM2, TM3* or *MKRS* (*oskar*), in order to detect zygotic effects on the incidence of atrophy among 3-7 day-old dysgenic F1 offspring. Offspring from the same cross are represented consecutively on the graph. While *bruno* alleles act as consistent dominant suppressors in both single and double mutants, *oskar* deficiencies and mRNA null mutants exhibit only stochastic effects on the atrophy phenotype, suggesting that the mechanism of *bruno* suppression of F1 atrophy is independent of *oskar* mRNA function. C-D) Representative germline development among atrophied and non atrophied ovaries of *bruno^QB^/+* and *CyO/+* dysgenic offspring (B). C) Atrophied ovaries lack any vasa-positive germline cells in the ovarioles, including GSCs. Nuclear Vasa external to Hts-1B1 corresponds to somatic ovarian sheath [35]. D) Non-atrophied (*i.e*. morphologically normal) ovaries exhibit the full range of developing oocytes, including GSCs (white arrow). Arrowheads correspond to cystoblasts (CBs), the undifferentiated daughters of GSCs. Representative ovaries in both C and D are *CyO/+*, however, atrophied ovaries (C) are more common among *bruno^QB^/+* (B). Cytological markers for C and D are hu li tai shao (Hts-1B1), which labels somatic follicle cell membranes the circular fusomes of GSCs and CBs, and the spectrosomes that connect cells in the developing cyst, Vasa, which labels cytoplasm of germline cells and nuclei of the ovarian sheath, and DAPI staining of nuclear DNA. The individual numerical values required to generate bar plots in panels A and B can be found in S9 Data and S10 Data, respectively.

Notably, the non-recessive, fertility enhancing effects of *bruno* loss-of-function alleles in dysgenic females contrasts their effects on the fertility of non-dysgenic females, where they act as recessive female steriles [49]. Our observations therefore suggest that a novel phenotype of *bruno* alleles is revealed by the dysgenic female germline. Furthermore, our observation that reduced *bruno* dosage suppresses ovarian atrophy is fully consistent with our observation that tolerant alleles are associated with reduced *bruno* expression (Fig 4D).

### *bruno’s* effects on ovarian atrophy are associated with germline stem cell retention

*bruno* is a translational regulator with three known functions in *Drosophila melanogaster* oogenesis. At the start of oogenesis in the ovarian substructure called the germaria, *bruno* is required to promote the differentiation of cystoblasts (CBs) the immediate daughters of germline stem cells (GSCs) [37,55]. In mid-oogenesis, Bruno protein blocks vitellogenesis if not properly sequestered by its mRNA target *oskar* [38,39]. Finally, in late oogenesis, Bruno repression of *oskar* translation is required to establish dorsovental (DV) patterning in the subsequent embryo [36]. This final role of *bruno* affects only the morphology of egg chambers, and but not their production, suggesting that it can not account for *bruno’s* effects in dysgenic germlines. We therefore focused on *bruno’s* earlier roles in GSC differentiation and vitellogenesis, which are distinguishable by their dependency on *oskar* mRNA. While *bruno’s* functions in GSC differentiation are independent of *oskar* mRNA [36,37,56], *bruno* and *oskar*’s impact on vitellogenesis are interdependent, due to the requirement for *oskar* mRNA to sequester Bruno protein [39].

To determine if the effect of *bruno* alleles on hybrid dysgenesis are independent of *oskar* mRNA, we examined whether two *oskar* mRNA null alleles, *osk^0^* and *osk^A87^*, as well as an *oskar* deficiency, affected the incidence of ovarian atrophy when compared to a balancer control (Fig 5B). We observed that *osk^0^* and *Df(3R)osk* exhibited no effect on the atrophy phenotype, while *osk^A87^* was associated with only a marginal increase in ovarian atrophy (*p* = 0.07). If *bruno* suppression of ovarian atrophy reflects reduced sequestration by *oskar* mRNA in the dysgenic female germline, atrophy should be enhanced by *oskar* mRNA mutants [57].

Comparison of single and double heterozygotes of *bruno* and *oskar*, also do not strongly suggest that *bruno* suppression is dependent on *oskar* mRNA dosage (Fig 5B). In comparisons involving two separate balancer 3 ^rd^ chromosomes, *Df(2L)bru/+; osk^A87^/+;* double heterozygotes did not differ from their single *Df(2L)bru/+;balancer/+heterozygous* siblings with respect to ovarian atrophy. A second double-heterozygote *Df(2L)bru/+;Df(3R)osk/+*; was associated with significantly increased atrophy when compared to the single heterozygote *(Df(2L)bru/+; balancer/+*). However, because this behavior was unique to the deficiency chromosome, and was not also exhibited by the *osk^A87^* mRNA null mutant that specifically eliminates *oskar* function [38], we suspect it is a synthetic consequence of hemizygosity in both deficiency regions, rather than evidence for a genetic interaction between *oskar* and *bruno* with respect to hybrid dysgenesis. Consistent with this, we also did not detect any Bruno mislocalization in the developing egg chambers of wild-type dysgenic females (S1 Fig), nor did we see any evidence of an arrest in mid-stage oogenesis in wild-type dysgenic ovaries, as occurs when Bruno is not sequestered by *oskar* mRNA [38,39].

To evaluate whether *bruno* suppression of ovarian atrophy could be explained by its oskar-independent role in GSC differentiation [55,58], we directly examined and compared GSCs between *bruno^QB^/+* and *CyO/+* dysgenic ovaries (Fig 5C, 5D). While we did not observe any direct evidence of delayed GSC differentiation in *bruno^QB^/+* germaria, we did observe that ovarioles containing developing egg chambers overwhelmingly retained all oogenic stages, including GSCs (Fig 5D). In contrast, atrophied ovaries lacked any developing egg chambers including GSCs (Fig 5C). These observations are consistent with recent evidence that GSC retention is a key determinant of *P*-element induced ovarian atrophy [35]. They further suggest that reduced *bruno* signaling for differentiation may stabilize GSCs in their niche, allowing them to be retained despite the genotoxic effect of *P*-elements.

## DISCUSSION

While the evolution of TE resistance through small-RNA-mediated silencing is a topic of immense research interest, the existence and evolution of tolerance factors that may reduce the fitness costs of TEs on the host remain undocumented. By opportunistically employing a panel of genotyped recombinant inbred lines, which uniformly lack small-RNA-mediated silencing of the recently invaded *P*-element, we have here uncovered the first example of segregating variation in host-TE tolerance in any organism. The natural variation in tolerance that we uncovered is unlikely to be exceptional. A complementary QTL analysis of tolerance in DSPR mapping population B has revealed a distinct peak of major effect in a different genomic location (Lama and Kelleher, *unpublished*). Furthermore, major QTL peak that we identified here in population A explains only about 35% of the heritable variation in tolerance. Therefore, other segregating variants in the population A RILs must also affect female germline response to *P*-element activity. Our inability to map these variants likely reflects the fact that they are rare, exhibit small effects, or both [30]. Finally, because our phenotyping scheme involved crossing genetically variable RILs to the same paternal strain, zygotic recessive alleles that are not found in the paternal Harwich genotype would remain undetected.

While differences in tolerance may be masked by resistance after small-RNA-mediated silencing evolves, our study reveals that in the absence of silencing, tolerance can be a major determinant of fitness. While the major peak we identified is modest in its functional effects, explaining ~14% of variation in ovarian atrophy, variation in fertility of this scale would be dramatic in the eyes of natural selection. Segregating tolerance alleles, such as the one we have detected here, could therefore be subject to strong positive selection during genome invasion by a novel TE. Tolerance may therefore be an important feature of the host evolutionary response to invading TEs. Indeed, the correlation between *P*-element dosage and the severity of hybrid dysgenesis is poor at best, causing many to suggest that other genetic factors, such as tolerance alleles, may also be important determinants of the dysgenic phenotype [59–61]. Furthermore, the hybrid dysgenesis induced by recent collections of *D. melanogster* tends to be mild when compared to collections from the 1970’s and 1980’s, providing circumstantial evidence that tolerance factors may have increased in frequency over time [61]. Once the causative variant responsible for the tolerance phenotype we uncovered here is identified, we will be poised to ask whether its increase in frequency has enhanced tolerance in extant populations.

Based on its high expression in the *Drosophila* female ovary [45], the presence of 22 SNPs that are in-phase with founder QTL alleles, its differential expression between tolerant and sensitive alleles, and the dominant suppressive effect of classical loss-of-function alleles on dysgenic ovarian atrophy, *bruno* is a very strong candidate for the source of causative variation in *P*-element tolerance that we mapped on chromosome 2L. Identifying the causative variant(s) within the very large (138 Kb) *bruno* locus, and understanding how its altered function relates to hybrid dysgenesis presents an exciting challenge for future work. On the surface, it is not obvious how *bruno* function could be related to *P*-element activity. Because Bruno physically interacts with the piRNA-pathway component Vasa [62] and localizes to nuage [63], the multifunctional germline organelle in which piRNA biogenesis occurs [reviewed in,64,65], a straightforward explanation is that *bruno* function is unrelated to tolerance, but rather suppresses piRNA-mediated resistance of *P*-elements. However, resistance suppression is inconsistent with several important aspects of piRNA biology and *bruno* function. First, piRNA-mediated silencing of *P*-elements is short-circuited in the absence of complementary maternally deposited piRNAs (absent from the RILs), and *P*-element derived piRNAs are exceptionally rare in the ovaries of young dysgenic females [42,66]. Thus, the dramatic suppression of ovarian atrophy exhibited by *bruno* alleles in young dysgenic females (Fig 5B), is developmentally inconsistent with piRNA-mediated silencing, which can occur only in older female offspring of dysgenic crosses [42]. Additionally, germline knock-down of *bruno* does not significantly affect TE expression, and if anything, is associated with increased expression of some TE families [67]. If *bruno* suppressed piRNA-silencing, reduced TE expression would be predicted upon knock-down.

We propose that our results are best explained *bruno*’s function in promoting GSC differentiation [37,55], which could determine the tolerance of GSCs to DNA damage resulting from *P*-activity. GSC maintenance is dependent on a balance between self-renewal and differentiation [reviewed in,68], and is disrupted the presence of DNA damage, leading to GSC loss [69]. We recently have discovered that the DNA damage response factor p53 is ectopically activated in the GSCs and CBs of dysgenic germlines [35], which explains why GSCs are frequently absent from dysgenic ovaries (Fig 5C, [15,34,35]). *bruno* loss-of-function alleles could therefore stabilize damaged GSCs in their niche in dysgenic germaria by reducing signals for differentiation. Indeed, loss-of-function mutations in two other GSC differentiation factors, *bam* and *bgcn*, have been associated with enhanced retention of GSCs in the niche of non-dysgenic germaria [70]. This model is fully consistent with our observation that *bruno* suppression of ovarian atrophy is accompanied by a rescue of oogenesis at all stages, including enhanced maintenance of GSCs (Fig 5C).

Our observations with *bruno* suggest an unexpected and novel role for developmental regulators of gametogenesis as determinants of germline tolerance of transposition. Interestingly, multiple regulators of female GSC maintenance and differentiation in *Drosophila melanogaster* exhibit recent or ongoing signatures of positive selection [71–73]. Tolerance to the selfish bacterial endosymbiont *Wolbacchia* has already been implicated in driving some of this adaptive evolution [74]. The fact that *bruno* alleles act as strong repressors of *P*-element hybrid dysgenesis suggests that another class of parasites, transposable elements, may also contribute to adaptive evolution of stem cell determinants.

## MATERIALS and METHODS

### Drosophila Strains and Husbandry

Recombinant inbred lines (RILs) from Population A were generously provided by Stuart Macdonald. Harwich (#4264), *Ced-12^c06760^/CyO* (#17781), *Rab6^GE13031^/CyO* (#26898), *Rab6^D23D^/CyO* (#5821), and *ThrRS^K04203^/CyO* (#10539), were obtained from the Bloomington *Drosophila* stock center. Harwich was sib-mated for one generation to increase homozygosity. *Bruno* and *oskar* mutants and deficiencies, in single and double heterozygous combinations, were generously provided by Paul MacDonald. *Canton-S* was obtained from Richard Meisel. All flies were maintained in standard cornmeal media. All experimental flies were maintained at 25°C.

### QTL Phenotyping

Virgin RIL females were crossed to Harwich males and flipped onto fresh food every 3-5 days.

Resulting F1 offspring were maintained for 3 days or 21 days at which point their ovaries were examined using a squash prep [60]. 21 day-old females were transferred onto new food every 5 days as they aged to avoid bacterial growth. For the squash prep, individual females were squashed in a food-dye solution allowed to incubate for ≥ 5 minutes. After incubation, the slide was examined for the presence of stage 14, chorionated egg chambers, which preferentially absorb the dye. In the interest of throughput, we assayed F1 females for the presence or absence of mature egg chambers: females who produced ≥ 1 egg chambers were scored as having non-atrophied ovaries, and females producing 0 egg chambers were scored as having atrophied ovaries. A phenotyping schematic is provided in Fig 1A.

Crosses and phenotyping were performed for 656 RILs across 24 experimental blocks for 3 day-old F1 females, and 606 RILs across 21 experimental blocks for 21 day-old F1 females. If fewer than 21 F1 offspring were phenotyped for a given cross, it was discarded and repeated if possible. In total, we phenotyped >20 3-day old and 21 days-old F1 female offspring for 592 RILs and 492 RILs, respectively, and 660 RILs were assayed for at least one of the age groups.

### QTL mapping

For age-class specific QTL mapping (3-day and 21-day old), the arcsine transformed proportion of F1 females (S2 Fig) with atrophied ovaries produced by each RIL (S1-S2 Data) was used as the response variable in a random effects multiple regression model that included experimental block and undergraduate experimenter.

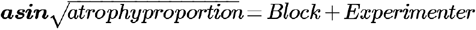

For the combined analysis of both age classes, we used the full set of arcsine transformed proportions, and accounted for female age as an additional fixed effect in the regression model.

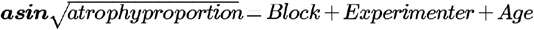

All models were fit using the lmer function from the lme4 package[75], and are described in S2 Table.

The raw residuals of the regression models above were used as the phenotypic response for QTL analysis (S3-S5 Data), implemented with the DSPRqtl package[29] in R 3.02 [76]. The output yields a <u>l</u>ogarithm of the <u>od</u>ds (LOD) score for the observed association between phenotype and genotype at 11,768 10 Kb intervals along the euchromatic arms of the X, 2 and 3^rd^ chromosomes. The LOD significance threshold was determined from 1,000 permutations of the observed data, and the confidence interval around each LOD peak was identified by a difference of −2 from the LOD peak position (Δ2-LOD), as well as a Bayesian credible interval [46].

### Broad sense heritability

Maternal genotype was added as a random effect to the models above, in order to determine the genetic variance in the phenotype (*V^G^*). *V^G^* was obtained by extracting the variance component for maternal genotype using the VarCorr() function from the nlme package [77]. Broad sense heritability (*H^2^*) was then estimated proportion of overall variance (*V^G^/V^P^*).

### Estimation of Founder Phenotypes and QTL phasing

To estimate the phenotypic effect of each founder at the QTL peak, we considered the residual phenotype for each used in QTL mapping and then determined the founder allele carried by the RIL at the LOD peak position [29]. RILs whose genotype at the LOD peak could not be assigned to a single founder with >0.95 probability were discarded.

Founder alleles were phased into phenotypic classes by identifying the minimal number of groups of required to describe phenotypic variation associated with the QTL peak [29]. Briefly, founder alleles were sorted based on their average estimated phenotypic effect, which was provided by the sampled RILs. Linear models containing all possible sequential partitions of founder alleles were then fit, and compared to a null model in which all founder alleles are in a single partition using an extra sum-of-squares *F*-test. The two-partition model with the highest *F*-statistic was retained and fixed only it if provided a significantly better fit *(P<* 10^−4^) than the null model. The two partitions of founder haplotypes were then fixed, and all possible three-partition models were explored. This process was continued until the model fit could not be improved.

### Identification of in-phase polymorphisms and TEs

Founders were assigned a “hard” genotype for all annotated TEs [78] and SNPs [29] in the QTL window if their genotype probability for a given allele was greater than 0.95 [29]. We then looked for alternate alleles (SNPs and TEs) that were in-phase with our inferred allelic classes [29,31]: the sensitive class (A3 and A5) and the tolerant class (A1, A2, A4, A6, and A7). A8 was excluded because its assignment to the sensitive or tolerant class differed between the data sets from 3-day old and 21-day old females.

### Identification of background matched, sensitive and tolerant RIL pairs

To identify RILs containing either the A4 (“tolerant”) or A5 (“sensitive”) haplotypes for the QTL window, we took advantage of the published, hidden-Markov model inferred genotypes for the population A RIL panel [29]. We first identified RILs that carried a contiguous A4 or A5 haplotype for the Δ2-LOD confidence interval for the combined analysis with a genotype probability of greater the 0.95 (Table 1). Then, for all possible RIL pairs (A4 and A5), we then calculated the number of 10 Kb genomic windows for which they carried the same RIL haplotype, also with a genotype probability of greater than 0.95. We selected three pairs of background matched RILs, which carry the same founder haplotype for 67% (11374 & 11120), 64% (11131 & 11200) and 60% (11435 & 11343) of genomic windows, but alternate haplotypes for the QTL window.

### F1 Fertility

Virgin female offspring of dysgenic crosses between tolerant (11120, 11200, 11343) and sensitive (11374, 11131, 11435) RILs and Harwich males were collected daily and placed individually in a vial with two *ywF10* males. Females were allowed to mate and oviposit for 5 days, and adults were discarded when the females reached 6 days of age. The presence and number of F2 offspring was quantified for each dysgenic female. The effect of genetic background and QTL haplotype on the presence and number of F1 offspring was assessed by logistic and linear regression models, respectively. Models were fit using the glm (logistic) and lm (linear) functions in R 3.02 [76].

### qPCR

Ovaries were dissected from young, 3 day old females from tolerant (11120, 11200, 11343) and sensitive (11374, 11131, 11435) RILs and homogenized in TRI-reagent (Sigma-Aldrich). RNA was extracted according to manufacturer instructions. Purified RNA was treated with DNAse, reverse transcribed using oligo(dT)15 primers and M-MLV RNAseH-, and then treated with RNAse H, according to manufacturer instructions (Promega). Synthesized cDNA was diluted 1:125 for qRT-PCR.

Abundance of *bruno* and *rp132* transcript was estimated using SYBR green PCR mastermix (Applied Biosystems) according to manufacturer instructions. Three biological replicates were evaluating for each genotype, with three technical measurements for each replicate, for a total of 9 measurements of each genotype. *Bruno* expression was estimated relative to *rp132* for each replicate according to a 5-point standard curve. Primers were as follows: *bruno*-F: 5’-CCCAGGATGCTTTGCATAAT-3’, *bruno*-R: 5’-ACGTCGTTCTCGTTCAGCTT-3’, *rp132*-F: CCGCTTCAAGGGACAGTATC, and *rp132*-F: GACAATCTCCTTGCGCTTCT. The relationship between genetic background and QTL haplotype with *bruno* expression was evaluated with mixed-effect linear regression, accounting for biological replicate as a random effect. The regression model was fit with the lme4 package [75] in R 3.02 [76].

### Mutational analysis of candidate genes

Single and double heterozygote mutant virgin females (*mutant/balancer*) were crossed to Harwich males at 25°C. Because the vast majority of *Drosophila* lab stocks are *P*-element free, with the exception of any *P*-element derived transgenes, these crosses are dysgenic. Resulting F1 dysgenic female offspring were collected and aged at 25°C for 3-7 days, when their ovaries were assayed using the squash prep described above [60]. The incidence of ovarian atrophy was then compared between *mutant/+* and *balancer/+* siblings from the same cross.

### Immunolabelling

Ovaries from 3-7 day old females offspring of dysgenic crosses were dissected and immediately fixed with 4% EM-grade methanol-free paraformaldehyde (Thermo Scientific). Ovaries were washed with 0.1% Triton X-100 in PBS, and blocked with 5% goat serum albumin (Sigma-Aldrich). Primary antibody concentrations were as follows: anti-Hts 1B1 1:4 (DSHB [79]), anti-Vasa 1:40 (DSHB), anti-Bruno 1:1000 (Provided by Paul MacDonald [50]), and anti-Orb 4H8 and 6H4 1:20 (DSHB [50]). Secondary antibody concentrations were 1:500.

### Microscopy

Ovaries were visualized with a SP8 Upright Confocal DM6000 CFS (Leica) Microscope, outfitted with a 60X oil immersion lens. Images were collected using an EM-CCD camera (Hamamatsu) and LAS-AF software (Leica).

## Acknowledgements

We are grateful to Stuart Macdonald, Paul MacDonald, and Richard Meisel for *Drosophila* strains, and Paul MacDonald for anti-Bruno antibodies. Ryan Castelluci, Ashka Shah and Wesley Adalume assisted in phenotyping atrophy among F1 offspring. Stuart Macdonald and Daniel Barbash, and three anonymous reviewers provided helpful discussion and feedback on this manuscript. This research was funded by NSF DEB #1457800 to Erin Kelleher.

---

**S1 Data. Proportion atrophy for 3 day-old F1 females**. For each sampled RIL, the strain number (matRIL), experimental block, student experimenter and proportion atrophy are provided.

**S2 Data. Proportion atrophy for 21 day-old F1 females**. For each sampled RIL, the strain number (matRIL), experimental block, student experimenter and proportion atrophy are provided.

**S3 Data. Residuals from combined regression, including both 3 day and 21 day-old females, used for QTL mapping**. For each sampled RIL, the strain number (matRIL), experimental block, student experimenter and residual are provided.

**S4 Data. Residuals from 3 day-old females, used for QTL mapping**. For each sampled RIL, the strain number (matRIL), experimental block, student experimenter and residual are provided.

**S5 Data. Residuals from 21 day-old females, used for QTL mapping**. For each sampled RIL, the strain number (matRIL), experimental block, student experimenter and residual are provided.

**S6 Data. Ovarian atrophy among dysgenic F1 offspring of background matched RILs**. Atrophy scores of 1 denote atrophied ovaries, while 0 indicates non-atrophied ovaries. For each female, the maternal RIL, background, and phenotype are provided.

**S7 Data. Fertility among dysgenic F1 offspring of background matched RILs**. For each female, the number of F2 offspring produced, the maternal RIL, background, and phenotype are provided.

**S8 Data. Bruno expression in background matched RILs**. *bruno* expression levels (relative to *rp132*), RIL genotype, background, phenotype and biological replicate are provided.

**S9 Data. Ovarian atrophy among F1 dysgenic offspring of candidate gene heterozygous mutant mothers**. The raw counts and proportion of atrophied and non-atrophied ovaries are provided for different offspring classes. Genotype indicates the zygotic genotype. Gene indicates gene affected by a heterozygous loss-of-function allele in the maternal genotype. Allele indicates the specific loss of function mutation.

**S10 Data. Ovarian atrophy among F1 dysgenic offspring of *bruno* and *oskar* mutant mothers**. The raw counts and proportion of atrophied and non-atrophied ovaries are provided for different offspring classes. Genotype indicates the zygotic genotype. Gene indicates whether the maternal genotype was heterozygous for a *bruno* loss-of-function allele, and *oskar* loss-of-function allele, or both. Allele indicates which mutations were found in the maternal genotype.

**S1 Table. In-phase polymorphisms within the QTL peak**. The chromosomal position on autosome 2L is provided for 36 inphase polymorphic SNPs. Coordinates are based on release 6 of the *D. melanogaster* genome [47]. REF and ALT indicate the nucleotide at this position found in the reference genome, or the alternative allele. The allele found in the founder genome is specificied as ref or alt (A1-A8). Putative effects on annotated genes are indicated according to the UCSC Genome Browser annotations [44].

**S2 Table. Random-effects ANOVA results for 3 day-old F1 females, 21-day F1 old females, and a combined analysis including both age classes**.

**S1 Fig. Bruno localization does not differ between dysgenic and non-dysgenic ovaries**. Bruno localization in mid-stage (4-6) oocytes of non-dysgenic (A) and dysgenic (B) females from reciprocal crosses between Canton-S and Harwich. In both genotypes, Bruno protein forms cytoplasmic, perinuclear rings.

**S2 Fig. Arcsine transformed and untransformed proportions of F1 atrophy among 3 day-old and 21-day old females**. Individual data points required to generate histograms is provided in S4 Data and S5 Data.

## Supporting Information

S6_Data.xlsx

S1_Data.xlsx

S2_Data.xlsx

S3_Data.xlsx

S5_Data.xlsx

S7_Data.xlsx

S9_Data.xlsx

S8_Data.xlsx

S4_Data.xlsx

S10 Data.xlsx

S1_Table.xlsx

S2_Table.xlsx

S2 Fig.tif

S1_Fig.tif

## References

1. Chénais B, Caruso A, Hiard S, Casse N. The impact of transposable elements on eukaryotic genomes: from genome size increase to genetic adaptation to stressful environments. Gene. 2012;509: 7–15. doi:10.1016/j.gene.2012.07.042

2. Vitte C, Panaud O. LTR retrotransposons and flowering plant genome size: emergence of the increase/decrease model. Cytogenet Genome Res. Karger Publishers; 2005;110: 91–107. doi:10.1159/000084941

3. Sun C, Shepard DB, Chong RA, López Arriaza J, Hall K, Castoe TA, et al. LTR retrotransposons contribute to genomic gigantism in plethodontid salamanders. Genome Biol Evol. 2012;4: 168–83. doi:10.1093/gbe/evr139

4. Kunarso G, Chia N-Y, Jeyakani J, Hwang C, Lu X, Chan Y-S, et al. Transposable elements have rewired the core regulatory network of human embryonic stem cells. Nat Genet. 2010;42: 631–4. doi:10.1038/ng.600

5. Lynch VJ, Leclerc RD, May G, Wagner GP. Transposon-mediated rewiring of gene regulatory networks contributed to the evolution of pregnancy in mammals. Nat Genet. 2011;43: 1154–9. doi:10.1038/ng.917

6. Startek M, Szafranski P, Gambin T, Campbell IM, Hixson P, Shaw CA, et al. Genome-wide analyses of LINE-LINE-mediated nonallelic homologous recombination. Nucleic Acids Res. 2015; doi:10.1093/nar/gku1394

7. Rajaei N, Chiruvella KK, Lin F, Astrom SU. Domesticated transposase Kat1 and its fossil imprints induce sexual differentiation in yeast. Proc Natl Acad Sci. 2014;111: 15491–15496. doi:10.1073/pnas.1406027111

8. Kapitonov V V, Jurka J. RAG1 Core and V(D)J Recombination Signal Sequences Were Derived from Transib Transposons. Nemazee D, editor. PLoS Biol. Public Library of Science; 2005;3: e181. doi:10.1371/journal.pbio.0030181

9. Silva-Sousa R, López-Panadès E, Casacuberta E. Drosophila telomeres: an example of co-evolution with transposable elements. Genome Dyn. 2012;7: 46–67. doi:10.1159/000337127

10. Aminetzach YT, Macpherson JM, Petrov DA. Pesticide resistance via transposition-mediated adaptive gene truncation in Drosophila. Science. 2005;309: 764–7. doi:10.1126/science.1112699

11. Mateo L, Ullastres A, González J. A Transposable Element Insertion Confers Xenobiotic Resistance in Drosophila. Feschotte C, editor. PLoS Genet. 2014;10: e1004560. doi:10.1371/journal.pgen.1004560

12. Hof AE van’t, Campagne P, Rigden DJ, Yung CJ, Lingley J, Quail MA, et al. The industrial melanism mutation in British peppered moths is a transposable element. Nature. Nature Research; 2016;534: 102–105. doi:10.1038/nature17951

13. Dupuy AJ, Fritz S, Largaespada DA. Transposition and gene disruption in the male germline of the mouse. Genesis. 2001;30: 82–8. Available: http://www.ncbi.nlm.nih.gov/pubmed/11416868

14. Spradling AC, Stern D, Beaton A, Rhem EJ, Laverty T, Mozden N, et al. The Berkeley Drosophila Genome Project gene disruption project: Single P-element insertions mutating 25% of vital Drosophila genes. Genetics. 1999;153: 135–77. Available: http://www.pubmedcentral.nih.gov/articlerender.fcgi?artid=1460730&tool=pmcentrez&rendertype=abstract

15. Dorogova N V., Bolobolova EU, Zakharenko LP. Cellular aspects of gonadal atrophy in Drosophila P-M hybrid dysgenesis. Dev Biol. 2017;424: 105–112. doi:10.1016/j.ydbio.2017.02.020

16. Noutsopoulos D, Markopoulos G, Vartholomatos G, Kolettas E, Kolaitis N, Tzavaras T. VL30 retrotransposition signals activation of a caspase-independent and p53-dependent death pathway associated with mitochondrial and lysosomal damage. Cell Res. 2010;20: 553–562. doi:10.1038/cr.2010.48

17. Roy BA, Kirchner JW. Evolutionary dynamics of pathogen resistance and tolerance. Evolution. 2000;54: 51–63. Available: http://www.ncbi.nlm.nih.gov/pubmed/10937183

18. Mauricio R. Natural selection and the joint evolution of toleranceand resistance as plant defenses. Evol Ecol. Kluwer Academic Publishers; 2000;14: 491–507. doi:10.1023/A:1010909829269

19. Blumenstiel JP. Evolutionary dynamics of transposable elements in a small RNA world. Trends Genet. Elsevier Ltd; 2011;27: 23–31. doi:10.1016/j.tig.2010.10.003

20. Yang P, Wang Y, Macfarlan TS. The Role of KRAB-ZFPs in Transposable Element Repression and Mammalian Evolution. Trends Genet. 2017; doi:10.1016/j.tig.2017.08.006

21. Senti K-A, Brennecke J. The piRNA pathway: a fly’s perspective on the guardian of the genome. Trends Genet. 2010;26: 499–509. doi:10.1016/j.tig.2010.08.007

22. Aravin AA, Hannon GJ, Brennecke J. The Piwi-piRNA pathway provides an adaptive defense in the transposon arms race. Science. 2007;318: 761–4. doi:10.1126/science.1146484

23. Schaack S, Gilbert C, Feschotte C. Promiscuous DNA: Horizontal transfer of transposable elements and why it matters for eukaryotic evolution. Trends Ecol Evol. 2010;25: 537–546. doi:10.1016/j.tree.2010.06.001

24. Kelleher ES. Reexamining the P-Element Invasion of Drosophila melanogaster Through the Lens of piRNA Silencing. Genetics. 2016;203: 1513–31. doi:10.1534/genetics.115.184119

25. Kofler R, Hill T, Nolte V, Betancourt AJ, Schlötterer C. The recent invasion of natural Drosophila simulans populations by the P-element. Proc Natl Acad Sci. 2015;112: 201500758. doi:10.1073/pnas.1500758112

26. Hill T, Schlötterer C, Betancourt AJ. Hybrid Dysgenesis in Drosophila simulans Associated with a Rapid Invasion of the P-Element. PLoS Genet. 2016;12: e1005920. doi:10.1371/journal.pgen.1005920

27. Kidwell MG. Evolution of hybrid dysgenesis determinants in Drosophila melanogaster. Proc Natl Acad Sci U S A. 1983;80: 1655–9. Available: http://www.pubmedcentral.nih.gov/articlerender.fcgi?artid=393661&tool=pmcentrez&rendertype=abstract

28. Kidwell MG, Frydrk T, Novy J. The hybrid dysgenesis potential of Drosophila melanogaster from diverse temporal and geographic origins. Drosoph Inf Serv. 1983;59: 63–69.

29. King EG, Merkes CM, McNeil CL, Hoofer SR, Sen S, Broman KW, et al. Genetic dissection of a model complex trait using the Drosophila Synthetic Population Resource. Genome Res. 2012;22: 1558–66. doi:10.1101/gr.134031.111

30. King EG, Macdonald SJ, Long AD. Properties and power of the Drosophila Synthetic Population Resource for the routine dissection of complex traits. Genetics. 2012;191: 935–49. doi:10.1534/genetics.112.138537

31. Long AD, Macdonald SJ, King EG. Dissecting complex traits using the Drosophila Synthetic Population Resource. Trends Genet. 2014;30: 488–495. doi:10.1016/j.tig.2014.07.009

32. Bucheton A. I transposable elements and I-R hybrid dysgenesis in Drosophila. Trends Genet. 1990;6: 16–21. Available: http://www.ncbi.nlm.nih.gov/pubmed/2158161

33. Malone CD, Lehmann R, Teixeira FK. The cellular basis of hybrid dysgenesis and Stellate regulation in Drosophila. Curr Opin Genet Dev. 2015;34: 88–94. doi:10.1016/j.gde.2015.09.003

34. Schaefer RE, Kidwell MG, Fausto-Sterling A. Hybrid Dysgenesis in Drosophila melanogaster: Morphological and Cytological Studies of Ovarian Dysgenesis. Genetics. 1979;92: 1141–52. Available: http://www.pubmedcentral.nih.gov/articlerender.fcgi?artid=1214061&tool=pmcentrez&rendertype=abstract

35. Tasnim S, Kelleher ES. p53 is required for female germline stem cell maintenance in P-element hybrid dysgenesis. Dev Biol. 2018;434: 215–220. doi:10.1016/j.ydbio.2017.12.021

36. Kim-Ha J, Kerr K, Macdonald PM. Translational regulation of oskar mRNA by bruno, an ovarian RNA-binding protein, is essential. Cell. 1995;81: 403–12. Available: http://www.ncbi.nlm.nih.gov/pubmed/7736592

37. Wang Z, Lin H. Sex-lethal is a target of Bruno-mediated translational repression in promoting the differentiation of stem cell progeny during Drosophila oogenesis. Dev Biol. 2007;302: 160–168. doi:10.1016/j.ydbio.2006.09.016

38. Jenny A, Hachet O, Závorszky P, Cyrklaff A, Weston MDJ, Johnston DS, et al. A translation-independent role of oskar RNA in early Drosophila oogenesis. Development. 2006;133. Available: http://dev.biologists.org/content/133/15/2827

39. Kanke M, Jambor H, Reich J, Marches B, Gstir R, Ryu YH, et al. oskar RNA plays multiple noncoding roles to support oogenesis and maintain integrity of the germline/soma distinction. RNA. 2015;21: 1096–109. doi:10.1261/rna.048298.114

40. Kidwell MG, Kidwell JF, Sved JA. Hybrid Dysgenesis in Drosophila melanogaster: A Syndrome of Aberrant Traits Including Mutation, Sterility and Male Recombination. Genetics. 1977;86: 813–33. Available: http://www.pubmedcentral.nih.gov/articlerender.fcgi?artid=1213713&tool=pmcentrez&rendertype=abstract

41. Engels WR, Preston CR. Hybrid dysgenesis in Drosophila melanogaster: the biology of female and male sterility. Genetics. 1979;92: 161–74. Available: http://www.pubmedcentral.nih.gov/articlerender.fcgi?artid=1213938&tool=pmcentrez&rendertype=abstract

42. Khurana JS, Wang J, Xu J, Koppetsch BS, Thomson TC, Nowosielska A, et al. Adaptation to P element transposon invasion in Drosophila melanogaster. Cell. 2011;147: 1551–63. doi:10.1016/j.cell.2011.11.042

43. Hartmann MA, Sekelsky J. The absence of crossovers on chromosome *4* in *Drosophila melanogaster*: Imperfection or interesting exception? Fly (Austin). Taylor & Francis; 2017;11: 253–259. doi:10.1080/19336934.2017.1321181

44. Casper J, Zweig AS, Villarreal C, Tyner C, Speir ML, Rosenbloom KR, et al. OUP accepted manuscript. Nucleic Acids Res. 2017;46: D762–D769. doi:10.1093/nar/gkx1020

45. Brown JB, Boley N, Eisman R, May GE, Stoiber MH, Duff MO, et al. Diversity and dynamics of the Drosophila transcriptome. Nature. 2014;512: 393–399. doi:10.1038/nature12962

46. Manichaikul A, Dupuis J, Sen S, Broman KW. Poor performance of bootstrap confidence intervals for the location of a quantitative trait locus. Genetics. Genetics Society of America; 2006;174: 481–9. doi:10.1534/genetics.106.061549

47. Hoskins RA, Carlson JW, Wan KH, Park S, Mendez I, Galle SE, et al. The Release 6 reference sequence of the Drosophila melanogaster genome. Genome Res. 2015;25: 445–58. doi:10.1101/gr.185579.114

48. King EG, Long AD. The Beavis Effect in Next-Generation Mapping Panels in Drosophila melanogaster. G3 (Bethesda). Genetics Society of America; 2017;7: 1643–1652. doi:10.1534/g3.117.041426

49. Schüpbach T, Wieschaus E. Female sterile mutations on the second chromosome of Drosophila melanogaster. II. Mutations blocking oogenesis or altering egg morphology. Genetics. 1991;129: 1119–36. Available: http://www.ncbi.nlm.nih.gov/pubmed/1783295

50. Filardo P, Ephrussi A. Bruno regulates gurken during Drosophila oogenesis. Mech Dev. 2003;120: 289–97. Available: http://www.ncbi.nlm.nih.gov/pubmed/12591598

51. Nilton A, Oshima K, Zare F, Byri S, Nannmark U, Nyberg KG, et al. Crooked, Coiled and Crimpled are three Ly6-like proteins required for proper localization of septate junction components. Development. 2010;137: 2427–2437. doi:10.1242/dev.052605

52. Chakraborty M, VanKuren NW, Zhao R, Zhang X, Kalsow S, Emerson JJ. Hidden genetic variation shapes the structure of functional elements in Drosophila. Nat Genet. Nature Publishing Group; 2018;50: 20–25. doi:10.1038/s41588-017-0010-y

53. Thibault ST, Singer MA, Miyazaki WY, Milash B, Dompe NA, Singh CM, et al. A complementary transposon tool kit for Drosophila melanogaster using P and piggyBac. Nat Genet. 2004;36: 283–287. doi:10.1038/ng1314

54. Parks AL, Cook KR, Belvin M, Dompe NA, Fawcett R, Huppert K, et al. Systematic generation of high-resolution deletion coverage of the Drosophila melanogaster genome. Nat Genet. 2004;36: 288–92. doi:10.1038/ng1312

55. Parisi MJ, Deng W, Wang Z, Lin H. The arrest gene is required for germline cyst formation during Drosophila oogenesis. Genesis. 2001;29: 196–209. Available: http://www.ncbi.nlm.nih.gov/pubmed/11309853

56. Kim-Ha J, Smith JL, Macdonald PM. oskar mRNA is localized to the posterior pole of the Drosophila oocyte. Cell. 1991;66: 23–35. Available: http://www.ncbi.nlm.nih.gov/pubmed/2070416

57. Lander ES, Linton LM, Birren B, Nusbaum C, Zody MC, Baldwin J, et al. Initial sequencing and analysis of the human genome. Nature. 2001;409: 860–921. doi:10.1038/35057062

58. Xin T, Xuan T, Tan J, Li M, Zhao G, Li M. The Drosophila putative histone acetyltransferase Enok maintains female germline stem cells through regulating Bruno and the niche. Dev Biol. 2013;384: 1–12. doi:10.1016/j.ydbio.2013.10.001

59. Bergman CM, Han S, Nelson MG, Bondarenko V, Kozeretska I. Genomic analysis of P elements in natural populations of Drosophila melanogaster. PeerJ. PeerJ Inc.; 2017;5: e3824. doi:10.7717/peerj.3824

60. Srivastav SP, Kelleher ES. Paternal Induction of Hybrid Dysgenesis in *Drosophila melanogaster* Is Weakly Correlated with Both *P*-Element and *hobo* Element Dosage. G3&#58; Genes|Genomes|Genetics. 2017; g3.117.040634. doi:10.1534/g3.117.040634

61. Ignatenko OM, Zakharenko LP, Dorogova N V, Fedorova SA. P elements and the determinants of hybrid dysgenesis have different dynamics of propagation in Drosophila melanogaster populations. Genetica. 2015;143: 751–9. doi:10.1007/s10709-015-9872-z

62. Webster PJ, Liang L, Berg CA, Lasko P, Macdonald PM. Translational repressor bruno plays multiple roles in development and is widely conserved. Genes Dev. Cold Spring Harbor Laboratory Press; 1997;11: 2510–21. Available: http://www.ncbi.nlm.nih.gov/pubmed/9334316

63. Snee MJ, Macdonald PM. Live imaging of nuage and polar granules: evidence against a precursor-product relationship and a novel role for Oskar in stabilization of polar granule components. J Cell Sci. 2004;117: 2109–20. doi:10.1242/jcs.01059

64. Kloc M, Jedrzejowska I, Tworzydlo W, Bilinski SM. Balbiani body, nuage and sponge bodies - The germ plasm pathway players. Arthropod Struct Dev. 2014;43: 341–348. doi:10.1016/j.asd.2013.12.003

65. Lehmann R. Germ Plasm Biogenesis—An Oskar-Centric Perspective. Current topics in developmental biology. 2016. pp. 679–707. doi:10.1016/bs.ctdb.2015.11.024

66. Brennecke J, Malone CD, Aravin AA, Sachidanandam R, Stark A, Hannon GJ. An epigenetic role for maternally inherited piRNAs in transposon silencing. Science. 2008;322: 1387–92. doi:10.1126/science.1165171

67. Czech B, Preall JB, McGinn J, Hannon GJ. A transcriptome-wide RNAi screen in the Drosophila ovary reveals factors of the germline piRNA pathway. Mol Cell. NIH Public Access; 2013;50: 749–61. doi:10.1016/j.molcel.2013.04.007

68. Sherman MH, Bassing CH, Teitell MA. Regulation of cell differentiation by the DNA damage response. Trends Cell Biol. NIH Public Access; 2011;21: 312–9. doi:10.1016/j.tcb.2011.01.004

69. Ma X, Han Y, Song X, Do T, Yang Z, Ni J, et al. DNA damage-induced Lok/CHK2 activation compromises germline stem cell self-renewal and lineage differentiation. Development. 2016;143: 4312–4323. doi:10.1242/dev.141069

70. Jin Z, Kirilly D, Weng C, Kawase E, Song X, Smith S, et al. Differentiation-Defective Stem Cells Outcompete Normal Stem Cells for Niche Occupancy in the Drosophila Ovary. Cell Stem Cell. 2008;2: 39–49. doi:10.1016/j.stem.2007.10.021

71. Civetta A, Rajakumar SA, Brouwers B, Bacik JP. Rapid evolution and gene-specific patterns of selection for three genes of spermatogenesis in Drosophila. Mol Biol Evol. 2006;23: 655–62. doi:10.1093/molbev/msj074

72. Bauer DuMont VL, Flores HA, Wright MH, Aquadro CF. Recurrent positive selection at bgcn, a key determinant of germ line differentiation, does not appear to be driven by simple coevolution with its partner protein bam. Mol Biol Evol. 2007;24: 182–91. doi:10.1093/molbev/msl141

73. Flores HA, DuMont VLB, Fatoo A, Hubbard D, Hijji M, Barbash DA, et al. Adaptive Evolution of Genes Involved in the Regulation of Germline Stem Cells in *Drosophila melanogaster* and *D. simulans*. G3&#58; Genes|Genomes|Genetics. 2015;5: 583–592. doi:10.1534/g3.114.015875

74. Flores HA, Bubnell JE, Aquadro CF, Barbash DA. The Drosophila bag of marbles Gene Interacts Genetically with Wolbachia and Shows Female-Specific Effects of Divergence. Malik HS, editor. PLOS Genet. 2015;11: e1005453. doi:10.1371/journal.pgen.1005453

75. Bates D, Mächler M, Bolker B, Walker S. Fitting Linear Mixed-Effects Models Using lme4. J Stat Softw. 2015;67: 1–48. doi:10.18637/jss.v067.i01

76. R Development Core Team. R: A Language and Environment for Statistical Computing [Internet]. Vienna, Austria; 2008. Available: http://www.r-project.org

77. Pinheiro J, Bates D, DebRoy S, Sarkar D, R Core Team. {nlme}: Linear and Nonlinear Mixed Effects Models [Internet]. 2016. Available: http://cran.r-project.org/package=nlme

78. Cridland JM, Macdonald SJ, Long AD, Thornton KR. Abundance and distribution of transposable elements in two Drosophila QTL mapping resources. Mol Biol Evol. 2013;30: 2311–27. doi:10.1093/molbev/mst129

79. Zaccai M, Lipshitz HD. Role ofAdducin-like (hu-li tai shao) mRNA and protein localization in regulating cytoskeletal structure and function duringDrosophila oogenesis and early embryogenesis. Dev Genet. 1996;19: 249–257. doi:10.1002/(SICI)1520-6408(1996)19:3<249::AID-DVG8>3.0.CO;2-9

